## Extended Data Figures for "*In Vivo* Metabolic Tagging and Targeting of Circulating Red Blood Cells"

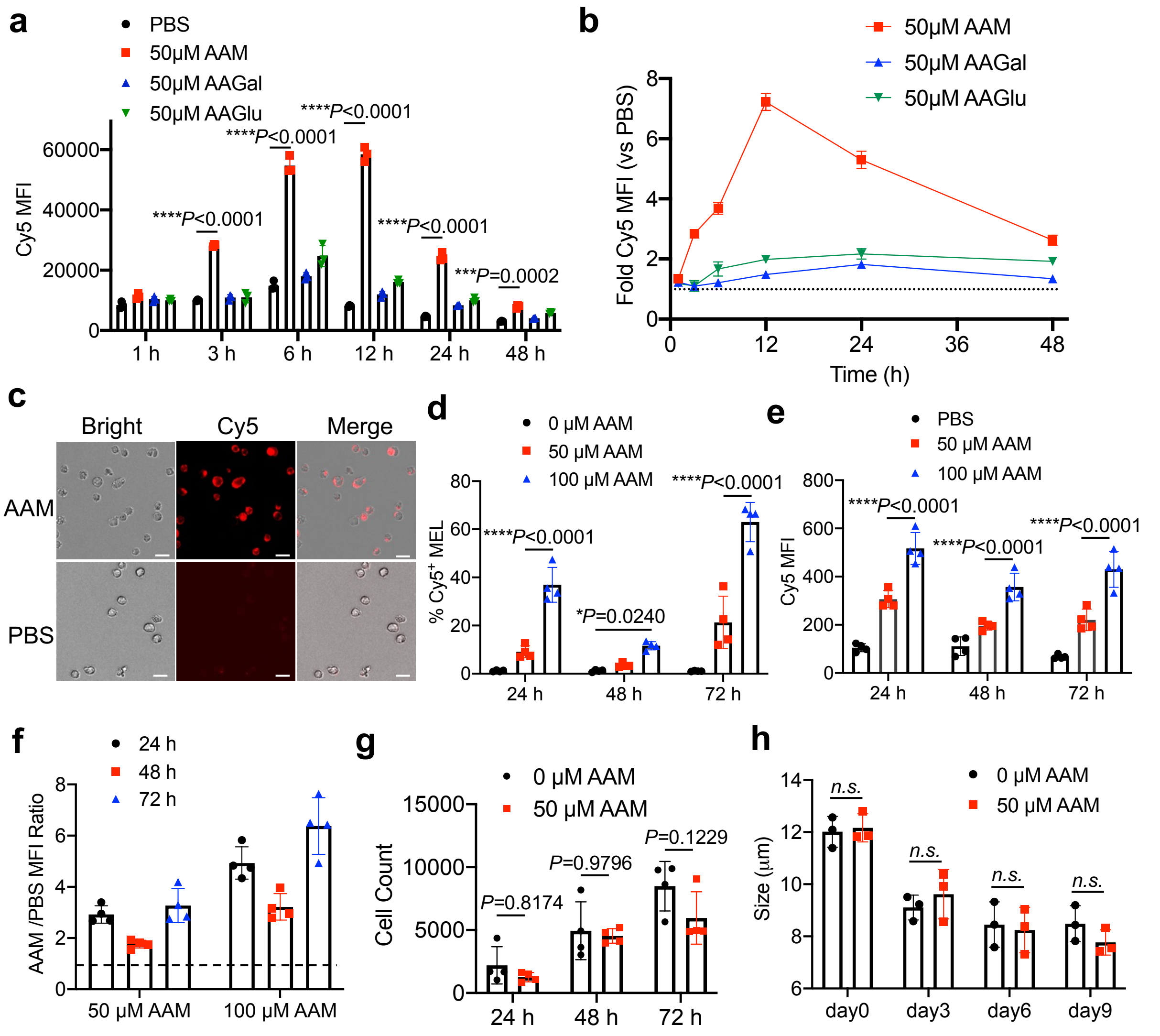


**Extended Data Figure 1. Metabolic glycan labeling of MEL cells.** (a) Mean Cy5 fluorescence intensity of MEL cells after incubation with azido-sugars or PBS for varied time (1, 3, 6, 12, 24, or 48 h) and staining with DBCO-Cy5 for 1 h. A concentration of 50 µM was used for all azido-sugars (AAM, AAGal, or AAGlu). (b) Cy5 fluorescence intensity ratio of MEL cells (azido-sugar/PBS) after incubation with azido-sugars for 1, 3, 6, 12, 24, and 48 h, respectively. (c) Fluorescence images of MEL cells treated with AAM for 24 h and then incubated with DBCO-Cy5 for 1 h. Scale bar: 10 µm. (d-g) MEL cells were treated with different concentrations of AAM for 24, 48 and 72 h, respectively, with fresh AAM added at 48 h. Shown are (d) % $Cy5^+$ MEL cells, (e) mean Cy5 fluorescence of MEL cells, (f) Cy5 fluorescence ratio of MEL cells (AAM/PBS), and (g) counts of MEL cells after azido-sugar treatment. (h) Size of cells during the differentiation process of MEL cells in the presence or absence of AAM. All the numerical data are presented as mean ± SD (one-way ANOVA with post hoc Fisher's LSD test was used; $0.01 < *P \leq 0.05$; $**P \leq 0.01$; $***P \leq 0.001$; $****P \leq 0.0001$).

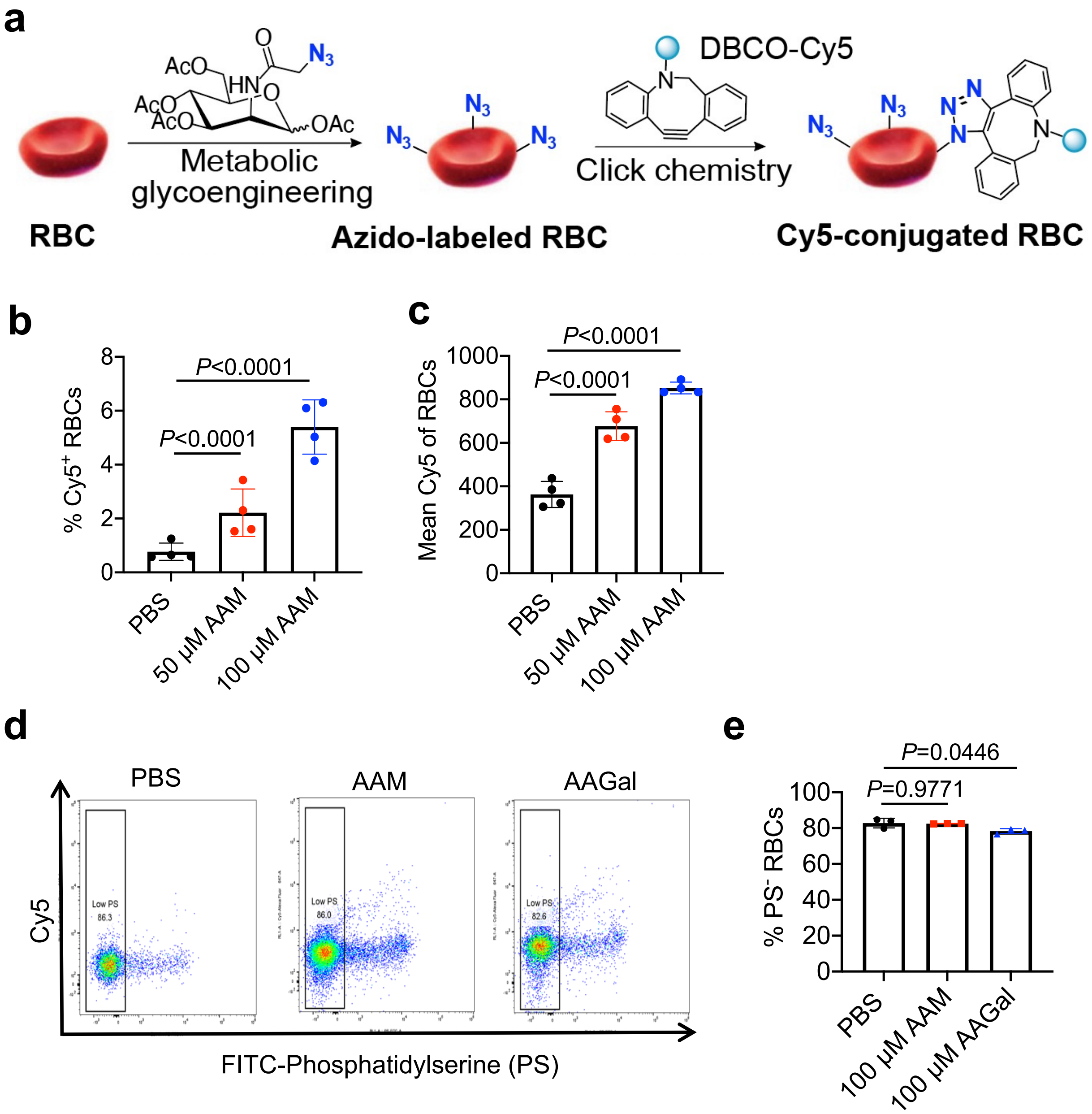


**Extended Data Figure 2. Metabolic glycan labeling of mouse RBCs *in vitro*.** (a) Schematic illustration of metabolic labeling of mouse RBCs with azido groups and subsequent conjugation of DBCO-molecules via click chemistry. (b) % $Cy5^+$ RBCs and (c) mean Cy5 fluorescence intensity of RBCs after 24-h incubation with AAM or PBS and 1-h staining with DBCO-Cy5. (d) Representative phosphatidylserine (PS) histogram of mouse RBCs after 24-h incubation with AAM or AAGal or PBS. (e) % $PS^+$ RBCs after 24-h incubation with AAM or AAGal or PBS. All the numerical data are presented as mean ± SD (one-way ANOVA with post hoc Fisher's LSD test was used; $0.01 < {*P} \leq 0.05$; $**P \leq 0.01$; $***P \leq 0.001$; $****P \leq 0.0001$).

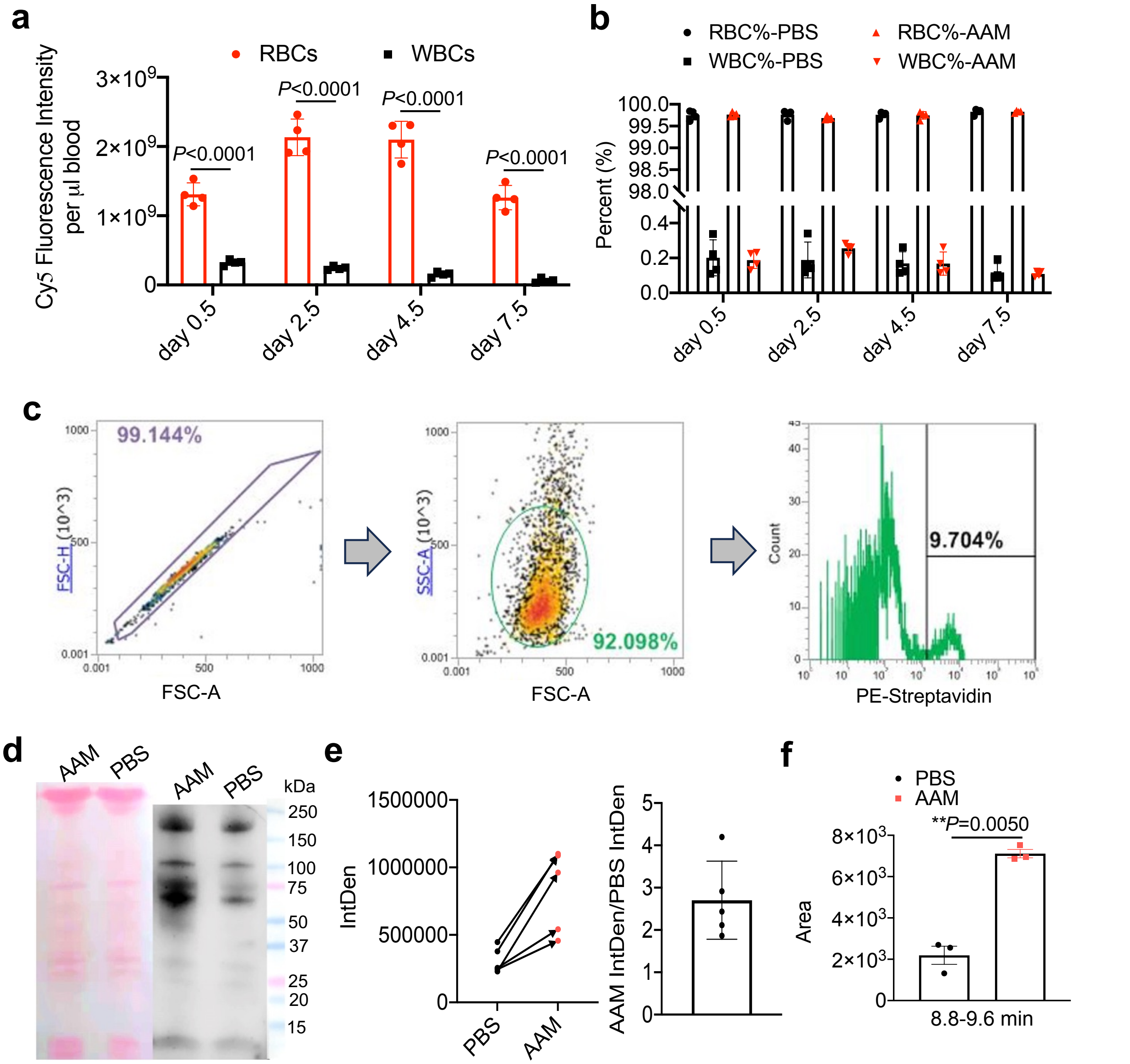


**Extended Data Figure 3. AAM can metabolically label glycoproteins and glycolipids of RBCs.** C57BL/6 mice were i.v. injected with AAM or PBS twice daily for three days. (a) Cy5 fluorescence intensity of RBCs and WBCs that were harvested at different times and stained with DBCO-Cy5. (b) % RBCs and WBCs in the blood at different times post injections of AAM or PBS. (c) Representative flow plots for analyzing azido-labeled RBCs. (d) Western blot analysis of RBCs isolated from mice at 14 days post AAM or PBS injections. Proteins extracted from RBCs were incubated with DBCO-biotin, run on a gel, and detected with streptavidin-HRP and chemiluminescence imaging. (e) Quantification of protein band signals in (d). (f) Quantification of Cy5-conjugated lipids from HPLC profiles (shown in Main Figure 2d). Lipids from RBCs were stained with DBCO-Cy5 and run on HPLC. All the numerical data are presented as mean ± SD (two-tailed Welch's t-test was used; $0.01 < {}^{*}P \leq 0.05$; $^{**}P \leq 0.01$; $^{***}P \leq 0.001$; $^{****}P \leq 0.0001$ ).

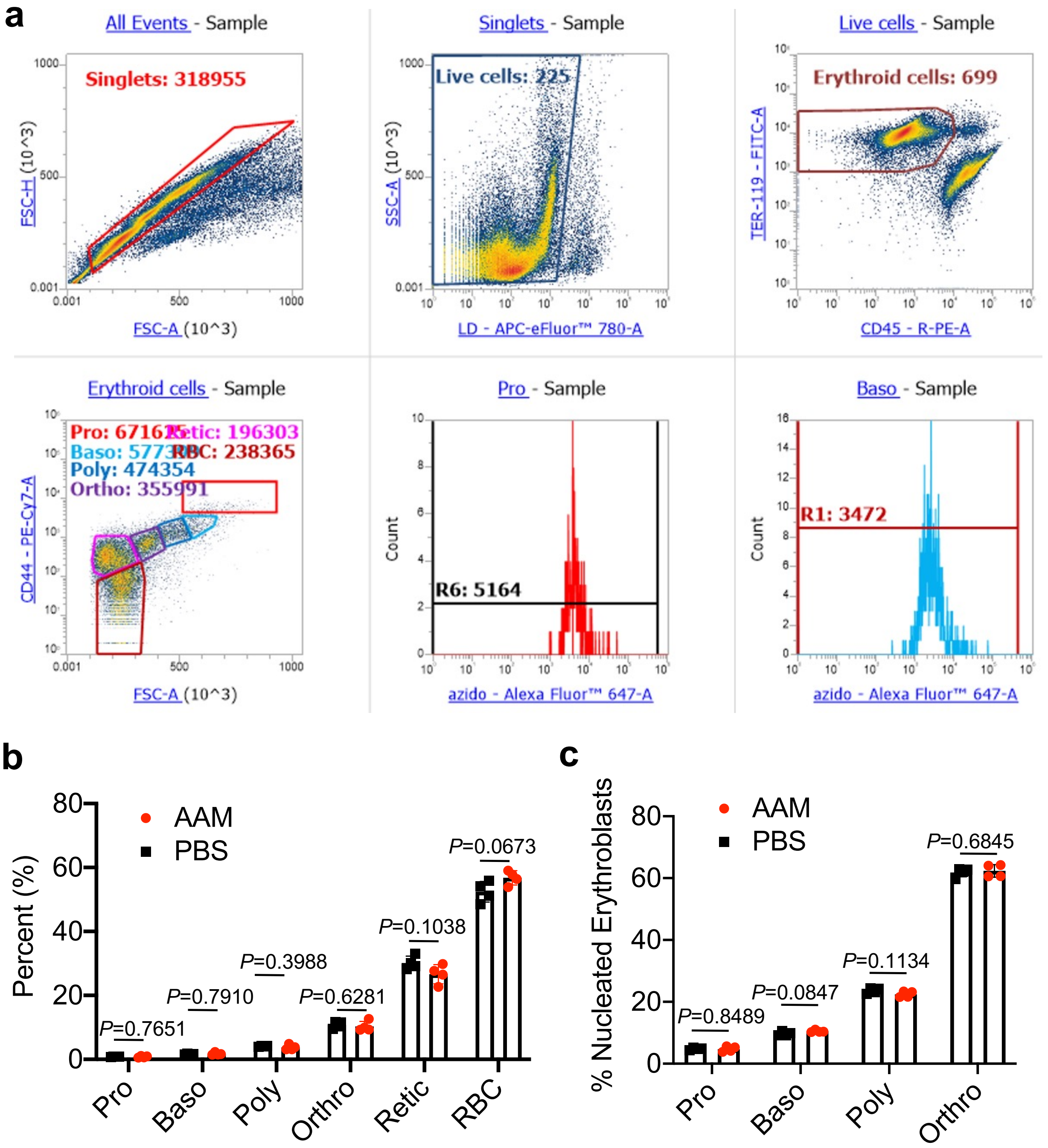


**Extended Data Figure 4. Intravenously injected AAM successfully labels RBC precursor cells in the bone marrow.** C57BL/6 mice were i.v. injected with AAM or PBS twice daily for three days. Bone marrow was harvested at 48 h post injections of AAM or PBS. (a) Representative flow plots for analyzing different erythroid lineage cells in the bone marrow. (b) Percentages of different RBC precursor cells in the bone marrow at 2 days post injections of AAM or PBS. (c) Percentages of nucleated erythroid lineage cells in the bone marrow at 2 days post injections of AAM or PBS. All the numerical data are presented as mean ± SD (one-way ANOVA with post hoc Fisher's LSD test was used; $0.01 < *P \leq 0.05$; $**P \leq 0.01$; $***P \leq 0.001$; $****P \leq 0.0001$).

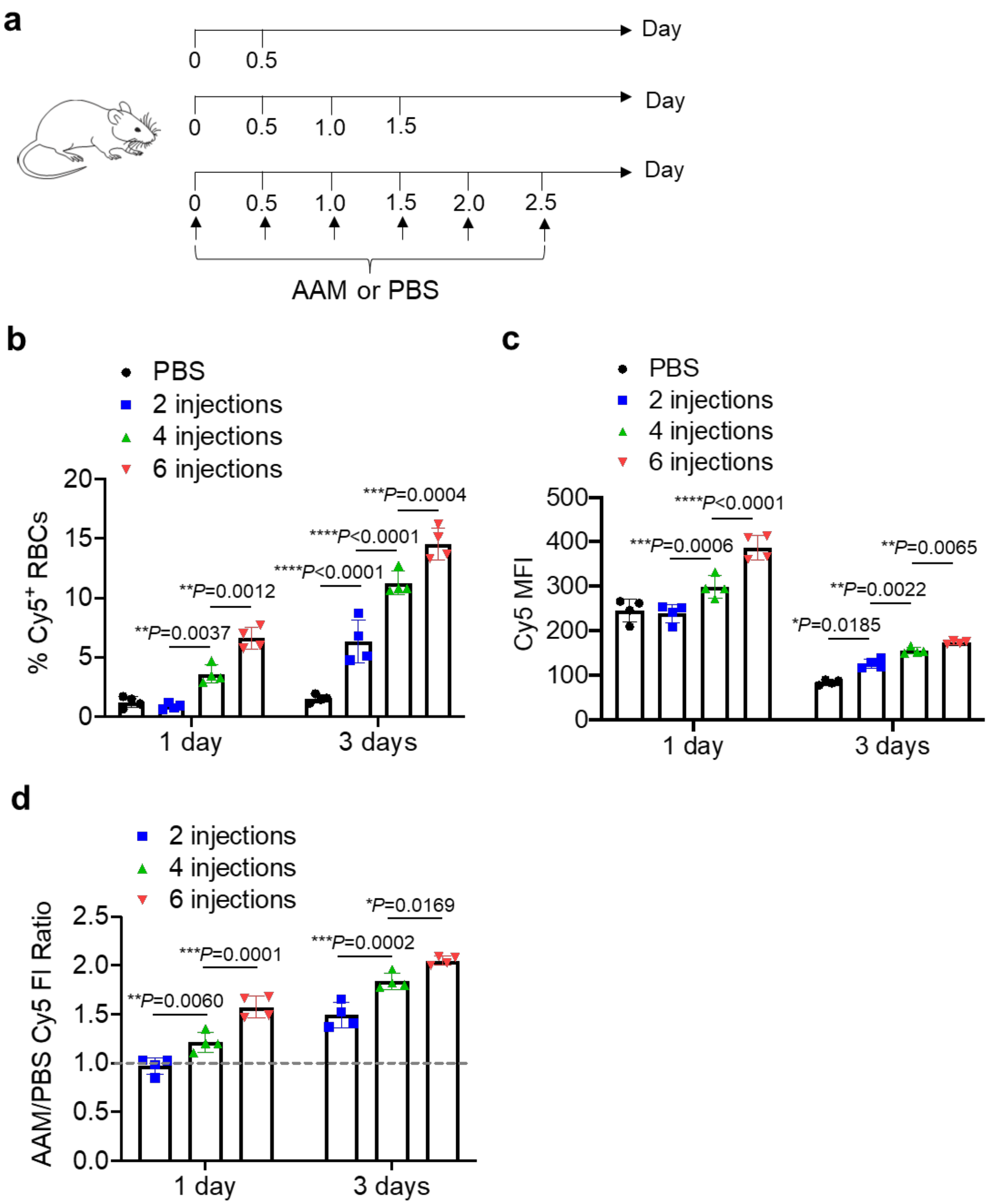


**Extended Data Figure 5. Intraperitoneally injected AAM metabolically labels RBCs with azido groups *in vivo* in a dose frequency dependent manner.** (a) Timeframe for the *in vivo* RBC labeling study. AAM (200 mg/kg) or PBS was intraperitoneally injected for 0, 2, 4, and 6 times, respectively with an interval of 12 h. RBCs were isolated at different times and incubated with DBCO-Cy5 for the detection of cell-surface azido groups. (b) Percentages of $Cy5^+$ RBCs at 1 or 3 days post the last injection of AAM or PBS. (c) Mean Cy5 fluorescence intensity of RBCs at 1 or 3 days post the last injection of AAM or PBS. (d) Cy5 fluorescence intensity ratio of RBCs (AAM/PBS) in (c). All the numerical data are presented as mean ± SD (one-way ANOVA with post hoc Fisher's LSD test was used; $0.01 < {*P} \leq 0.05$; $**P \leq 0.01$; $***P \leq 0.001$; $****P \leq 0.0001$).

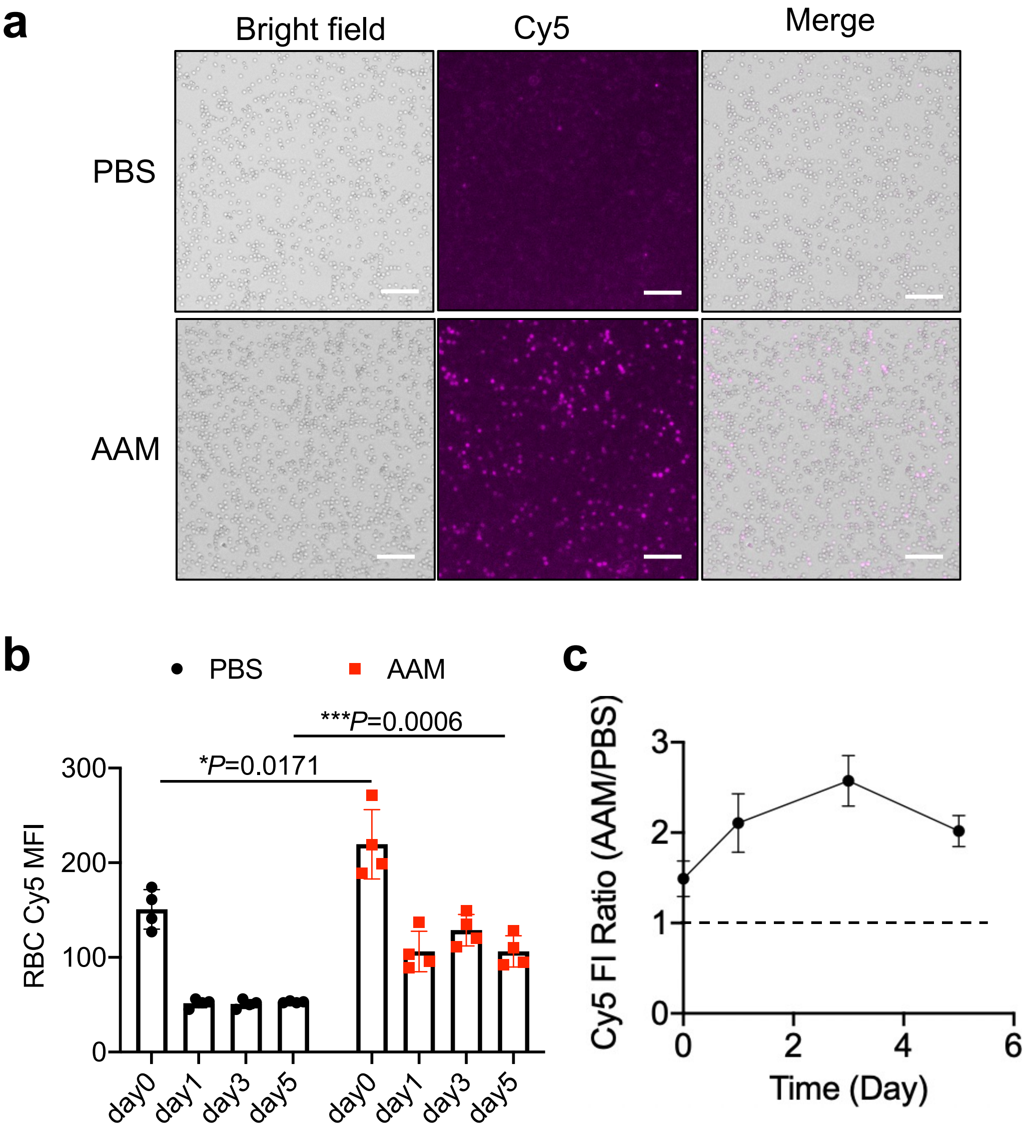


**Extended Data Figure 6. DBCO-Cy5 conjugated to azido-labeled RBCs shows stable membrane retention.** (a-c) AAM or PBS was i.v. injected into C57BL/6 mice twice daily for three days. After 7 days, RBCs were isolated and incubated with DBCO-Cy5 for 1 h *ex vivo*. After washing, RBCs were stored in the Alsever's solution for different times, prior to flow cytometry analysis. (a) Representative fluorescence images of RBCs isolated from AAM- or PBS-treated mice. Scale bar: 100 μm. (b) Cy5 fluorescence intensity of RBCs over time. (c) Cy5 fluorescence intensity ratio of RBCs (AAM/PBS) in (b). All the numerical data are presented as mean ± SD (one-way ANOVA with post hoc Fisher's LSD test was used; $0.01 < *P \leq 0.05$; $**P \leq 0.01$; $***P \leq 0.001$; $****P \leq 0.0001$).

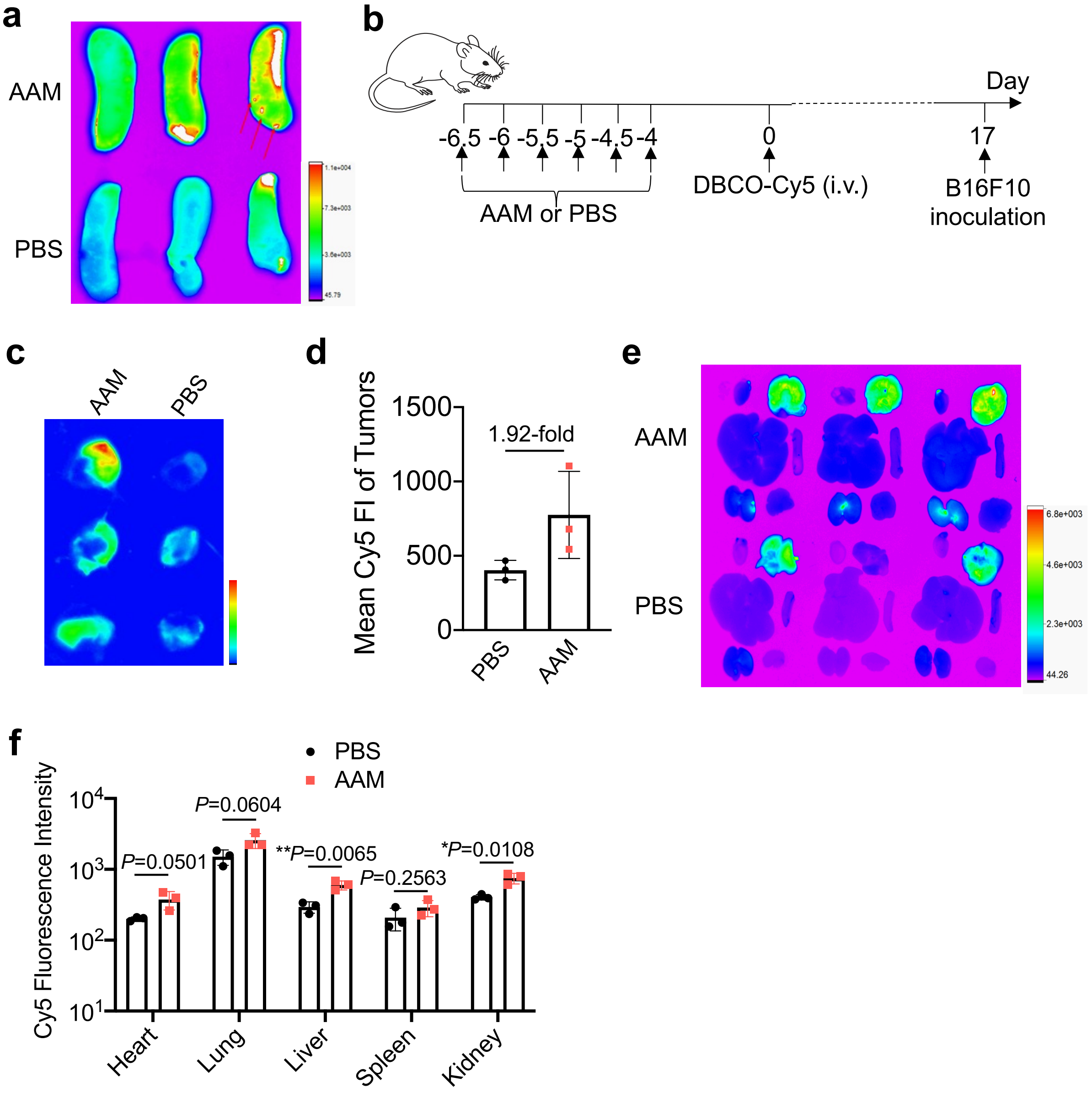


**Extended Data Figure 7.** ***In vivo*** **conjugation of DBCO-Cy5 onto RBCs enables fluorescence imaging of blood vessels and tissues.** (a) AAM or PBS was i.v. injected into Balb/c mice twice daily for three days (day -9.5 to day -7), followed by subcutaneous injection of 4T1 tumor cells on day -3 and i.p. injection of DBCO-Cy5 on day 0. Shown are IVIS images of spleens harvested from mice at 21 days post injection of DBCO-Cy5. The terminal arterioles structures in the spleen are indicated by the red arrows. (b-f) Timeframe for RBC labeling and B16F10 tumor imaging study. AAM or PBS was i.v. injected twice daily for three days, followed by i.v. injection of DBCO-Cy5 on day 0 and subcutaneous injection of B16F10 tumor cells on day 17. (c) IVIS imaging of B16F10 tumors at 7 days post tumor inoculation. (d) Quantified Cy5 fluorescence intensity of B16F10 tumors in (c). (e) IVIS imaging of major organs harvested from AAM- or PBS-treated mice on day 24. (f) Quantified Cy5 fluorescence intensity of organs from (e). All the numerical data are presented as mean ± SD (two-tailed Welch's t-test was used; $0.01 < *P \leq 0.05$; $**P \leq 0.01$; $***P \leq 0.001$; $****P \leq 0.0001$ ).

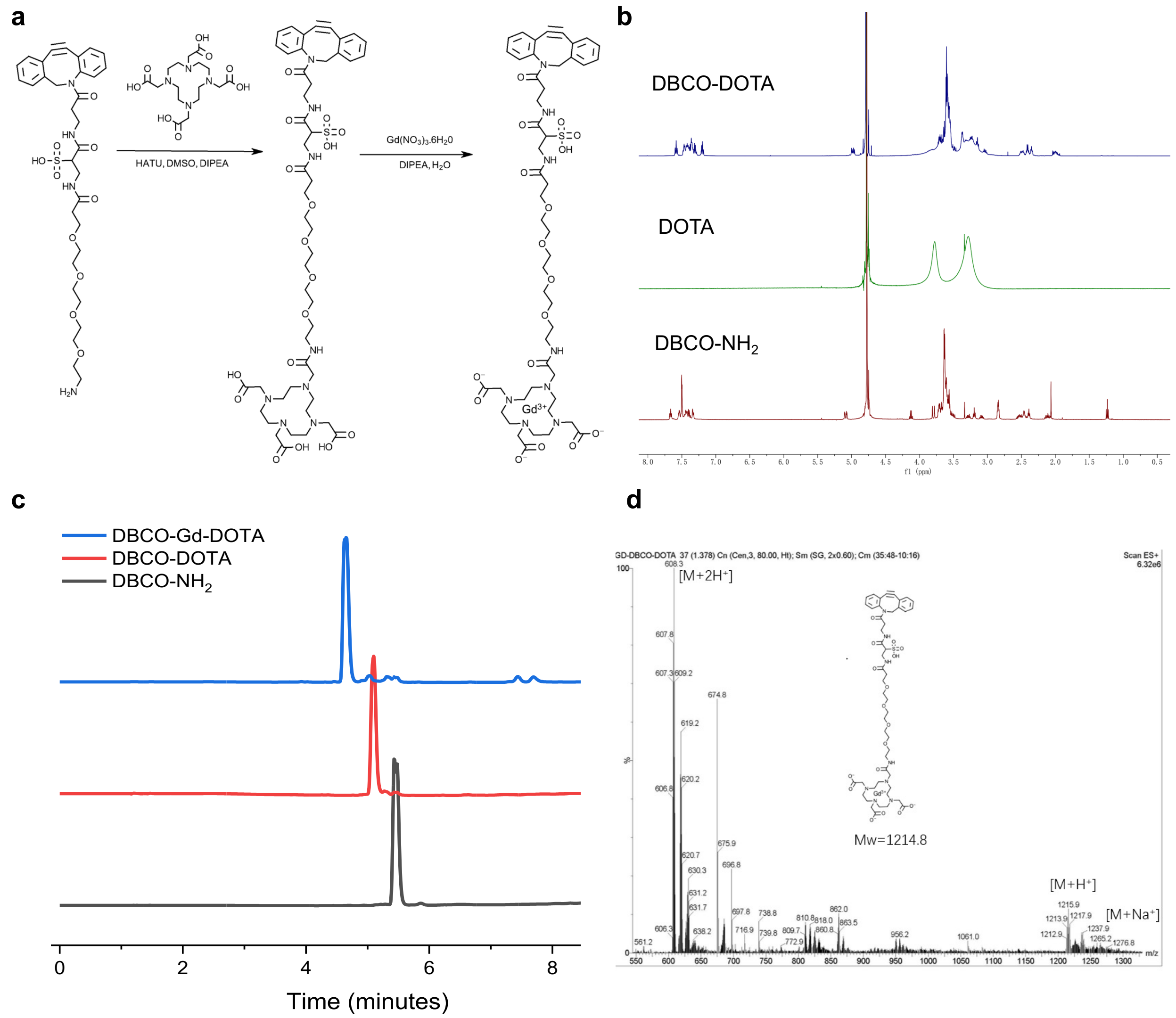


**Extended Data Figure 8. Synthesis and characterization of DBCO-DOTA-Gd.** (a) Synthesis route of DBCO-DOTA-Gd. (b) NMR spectra of $DBCO-NH_2$, DOTA, and DBCO-DOTA in $D_2O$. (c) HPLC profiles of $DBCO-NH_2$ (black), DBCO-DOTA (red), and DBCO-DOTA-Gd (blue). The detection wavelength was set at 310 nm. (d) Mass spectrum of DBCO-DOTA-Gd. The peaks for at $[M+H]^+$, $[M+2H]^{2+}$ and $[M+Na]^+$ can be found.

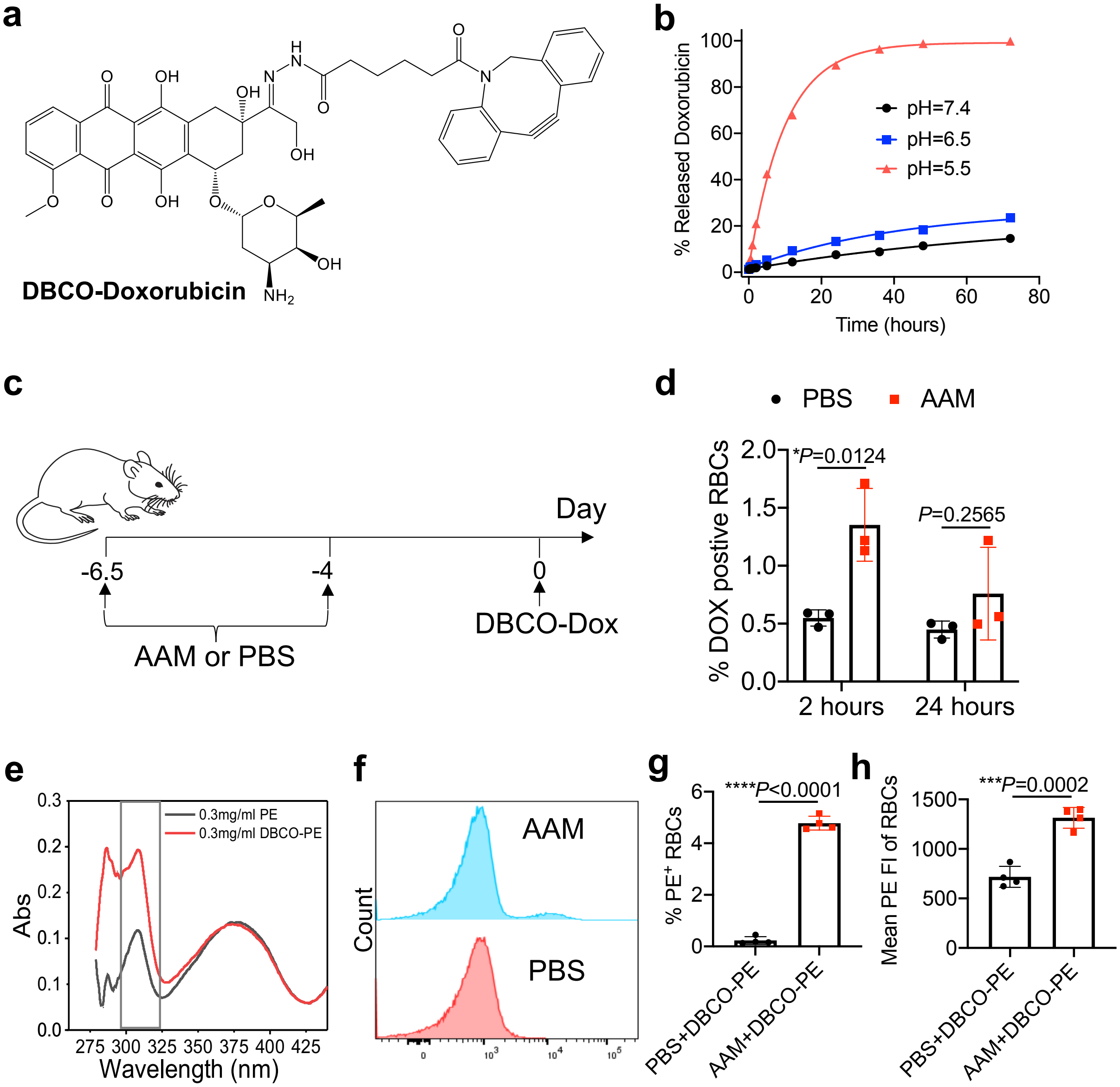


**Extended Data Figure 9. Azido-tagged RBCs enable conjugation of small-molecule and macromolecular cargos *in vitro* and *in vivo*.** (a) Synthesis of acid-labile DBCO-doxorubicin (DBCO-Dox). (b) Release kinetics of DBCO-Dox at different pH, as measured by HPLC. (c-d) C57BL/6 mice were i.v. injected with AAM twice daily for three days, and DBCO-Dox was i.v. injected at 4 days post the last AAM injection. The blood was collected after 2 or 24 h post DBCO-Dox injection. (c) Timeframe of study. (d) Percentages of Dox-containing RBCs at 2 or 24 h post DBCO-Dox injection. (e) UV spectra of PE and DBCO-functionalized PE. (f-h) Flow cytometry analysis of *in vitro* conjugation of PE to RBCs. RBCs were collected from AAM-treated mice at 14 days post AAM injection and incubated with DBCO-PE for 2 h. (f) Representative PE histogram of RBCs. (g) Percentages of PE-containing RBCs. (h) Mean PE fluorescence intensity of RBCs. All the numerical data are presented as mean ± SD (two-tailed Welch's t-test was used; $0.01 < *P \leq 0.05$; $**P \leq 0.01$; $***P \leq 0.001$; $****P \leq 0.0001$).

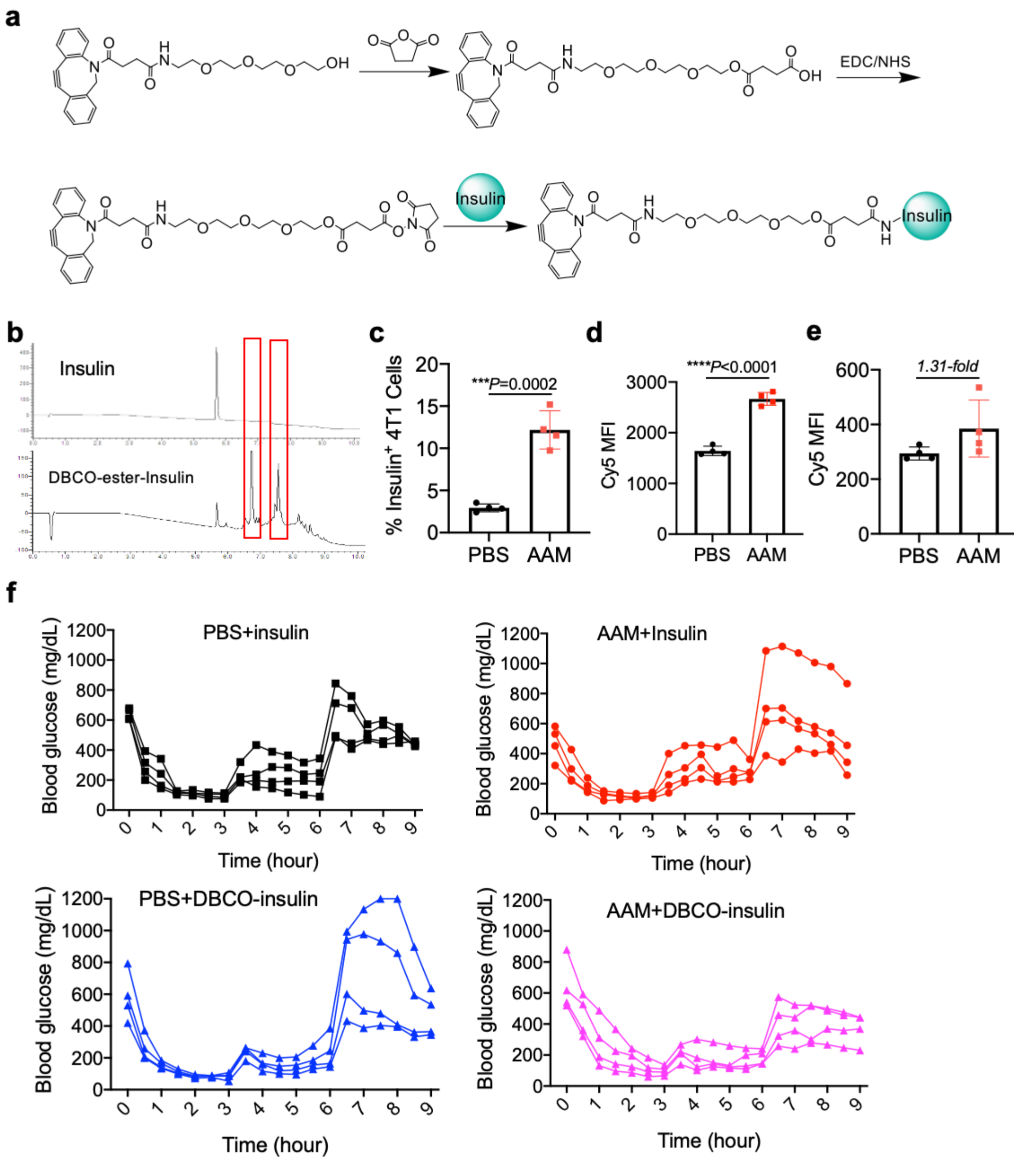


**Extended Data Figure 10. DBCO-insulin can conjugate to azido-labeled RBCs *in vivo* for improved blood glucose control.** (a) Synthesis of DBCO-insulin with an ester linkage. (b) HPLC profiles of insulin and DBCO-insulin with a detection wavelength of 280 nm. Peaks for DBCO-insulin are indicated. (c-d) 4T1 cells were treated with AAM or PBS for 24 h, followed by 1-h incubation with DBCO-insulin. Cell-surface insulin was detected by staining with rabbit anti-insulin and Cy5-conjugated goat anti-rabbit secondary antibody. Shown are (c) percentages of insulin-positive 4T1 cells and (d) Mean Cy5 fluorescence intensity of 4T1 cells. (e) Mean Cy5 fluorescence intensity of RBCs. C57BL/6 mice were i.v. injected with AAM or PBS twice a day for three days. After 14 days, RBCs were isolated and incubated with DBCO-insulin for 1 h. Cell-surface insulin was detected by staining with rabbit anti-insulin and Cy5-conjugated goat anti-Rabbit secondary antibody. (f) Blood glucose levels of individual mice for each group during the course of the glucose tolerance test. Mice were fasted for 12 h, and then 10 IU/kg DBCO-insulin or insulin was i.p. injected. Glucose was i.p. injected at 3 and 6 h. All the numerical data are presented as mean ± SD (two-tailed Welch's t-test was used; $0.01 < *P \leq 0.05$; $**P \leq 0.01$; $***P \leq 0.001$; $****P \leq 0.0001$).
